# A Simulation Analysis and Screening of Deleterious Non-Synonymous Single Nucleotide Polymorphisms (SNPs) in Human CDKN1A Gene

**DOI:** 10.1101/240820

**Authors:** G. M. Shazzad Hossain Prince, Trayee Dhar

## Abstract

CDKN1A also known as *p21^CIP1^ /p21^WAF1^*, a cyclin dependent kinase 1, interacts with proliferating cell nuclear antigen (PCNA) resulting in cell cycle inhibition in human. Non-synonymous single nucleotide polymorphisms (nsSNPs), which reside in the coding region of a gene, might distort the normal function of the corresponding protein. In silico analysis in this study followed many different algorithms. Following the final screening of 118 nsSNPs from dbSNP (NCBI), 12 missense SNPs (R19C (C→T), G23D (A→G), V25G (G→T), V25L (C→G), Q29P (A→C→G), F51L (C→T), E56K (A→G), T57I (C→T), G61R (C→G), G61D (A→G), Y151C (A→G) and R156W (C→G→T) were predicted to have deleterious effect by all the algorithms. Of them, R19C, G23D, F51L, Y151C and R156W occurred at the highly conserved site. G23D, F51L variants also occurred at the CDI domain. Homology structures of the protein predicted decrease of energy in mutant models. GV-GD scores predicted only two variants as neutral (V25L, F51L).

## 1. Introduction

CDKN1A or *p21^cip1^ /p21^WAF1^* negatively regulates cell cycle at G1 check point by binding to proliferating cell nuclear antigen (PCNA) which allows cells to repair damaged DNA. Cyclin dependent kinase inhibitor 1 or *p21^cip1^* inactivates PCNA. This promotes functions of *p53* (Yates et al., 2015). Dysfunction at G1 checkpoint may give rise to mutation and inappropriate cell proliferation which in turn might induce cancer progression (Bahl et al., 2000; Facher et al., 1997). CDKN1A or *p21^cip1^* also displays anti-apoptotic roles while inhibiting stress activated protein kinase (SAPK), apoptosis signal-regulating kinase 1 (ASK1), and inhibiting Fas-mediated apoptosis. (Yates et al., 2015). CDKN1A inhibits functional subunits of cyclinA-cdk2, cyclin E-cdk2, cyclin D-cdk4, and cyclin D-cdk6 and is unregulated by *TP53* pathway (Ravitz and Wenner, 1997). It plays central role in cellular growth arrest, terminal differentiation, and apoptosis. Direct regulation of p21 expression occurs by *p53*, and if *p21^cip1^ /p21^WAF1^* is cleaved by Caspase-3-mediated pathway, apoptosis of cancer cell follows (Ralhan et al., 2000). Mutations in *TP53* is found in association with chronic lymphocytic leukemia (CLL). Dysfunction in the interaction between *TP53* and CDKN1A can cause structural abnormality of *TP53* (Tracy et al., 2017).

Some common sequence variants, such as polymorphism on codon 31 in p21, was found associated with the procession of breast cancer. Mutation in CDKN1A, commonly known as or p21^cip1^ /p21^WAF1^, was hardly responsible for various types of cancers, despite being a principle downstream regulator of TP53 (Powell et al., 2002). But two common polymorphisms, C→A transversion at codon 31 (Ser→Arg) and C→T transition in the 3’UTR of exon 3, 20bp downstream from strop codon) in CDKN1A or p21^cip1^ /p21^WAF1^ has been suggested to induce development of some types of cancer i.e. esophageal cancer. (Bahl et al., 2000; Facher et al., 1997; Powell et al., 2002; Tracy et al., 2017). Another polymorphism A→G at codon 149 (Asp→Gly) was observed in esophageal squamous cell carcinomas (ESCCs) in Indian patients and was predicted to interfere with *p53* pathway as well as esophageal tumorigenesis (Bahl et al., 2000; Powell et al., 2002). Polymorphism in CDKN1A was also reported as a risk factor to Alzheimer’s disease (AD) (Yates et al., 2015). In silico methods with great accuracy was also followed to screen disease associated nsSNPs in many studies using computational tool such as-PolyPhen 2.0, SIFT, PANTHER, I-mutant 3.0, PhD-SNP, SNP&GO, Pmut, and Mutpred etc. (Ali et al., 2017; Kamaraj and Purohit, 2013; Mathe et al., 2006; Stojiljkovic et al., 2016). Transactivation activity of 1514 missense substitution was analyzed by a Align GV-GD algorithm which scores substitution from C0 (neutral) to C65 (deleterious) (Ali et al., 2017; Mathe et al., 2006). Conservation analysis, based on the evolutionary information, of a protein unfolds its amino acid positions along the chain, which is essential to retain its structural integrity and function (Ashkenazy et al., 2016). Many different computer based algorithms to predict missense variants should be optimized as well as sequence alignment to find the best result (Hicks et al., 2011). Application of many computational analysis was employed in this study to find deleterious nsSNPs of CDKN1A as well as their functional effect on the corresponding protein.

## 2. Methods

### 2.1. Collecting SNPs and Protein’s Sequence from the databases

Human CDKN1A gene SNPs were collected from the National Center for Biological Information (NCBI) online dbSNP database (https://www.ncbi.nlm.nih.gov/snp). From this server, the non-synonymous SNP (nsSNP) information (Chromosome coordinates and alleles) was retrieved for the computational analysis in this study (Sherry et al., 2001). Protein IDs from the analysis of nsSNPs in SIFT server (http://sift.jcvi.org/) were used to retrieve FASTA sequence of the protein encoded by Human CDKN1A gene from the Ensemble database (https://ensembl.org/). Ensemble is a growing database which has been working for aggregating, processing, integrating and redistributing genomic datasets (Zerbino et al., 2017).

### 2.2. Deleterious SNPs predicted by SIFT and Polyphen-2

Sorting intolerant from the tolerant (SIFT) server was used to predict the damaging nsSNPs (Kumar et al., 2009). It receives a query sequence and employs multiple alignment to predict either tolerated or damaging substitution in each position of the query sequence (Ng and Henikoff, 2003). nsSNPs with a normalized probability score less than 0.05 is predicted to be harmful and a score greater than 0.05 is considered as tolerated (Ng and Henikoff, 2006). Polyphen-2 server available at (http://genetics.bwh.harvard.edu/pph2/) was utilized, using the data gathered from SIFT result of the nsSNPs, to find either deleterious or neutral amino acid substitutions. By naive Bayesian classifier, it identifies the potential amino acid substitution and the effect of mutation (Adzhubei et al., 2010). It classifies SNPs as “Probably damaging” (0.85 to 1.00), “Possibly damaging’ (0.15 to 0.84) and “Benign” (<0.14) (Ali et al., 2017).

### 2.3. Disease related amino acid substitution by Predict-SNP, MAPP, PhD-SNP, and SNAP

Predict SNP, a consensus classifiers for prediction of disease related amino acid mutation, available at (https://loschmidt.chemi.muni.cz/predictsnp1/) with integrated tools-MAPP, PhD-SNP 2.06 and SNAP 1.1.30, was used to determine the deleterious or neutral nsSNPs (Bendl et al., 2014). Multivariate Analysis of Protein Polymorphism (MAPP) defines deleterious nsSNPs based on the physicochemical variation present in a column of protein sequence akignment (Stone and Sidow, 2005). Predictor of Human Deleterious Single Nucleotide Polymorphism (PhD-SNP) is a Support Vector Machine (SVM) which uses evolutionary information to sort out SNPs related to Mendelian and complex diseases from the neutral ones (Capriotti et al., 2006). SNAP screens out non-acceptable polymorphisms using neural network. It determines 80% of the deleterious substitution at 77% accuracy and 76% of the neutral substitutions at 80% accuracy (Bromberg and Rost, 2007).

### 2.4. Analysis of protein’s stability change upon amino acid substitution

#### 2.4.1. I-Mutant 3.0 server

A Support Vector Machine (SVM) based predictor I-Mutant 3.0, available at (http://gpcr2.biocomp.unibo.it/cgi/predictors/I-Mutant3.0/I-Mutant3.0.cgi, was used in this study to calculate the change in the stability of protein resulting from amino acid substitution (Capriotti et al., 2005). It computes the DDG (kcal/mol) value and RI value (reliability index) of a submitted mutation of a protein.

#### 2.4.2. MUpro server

MUpro server uses both the Support Vector Machine (SVM) and neural network to predict the change in the stability of the protein upon single site mutation. It predicts, with an 84% accuracy, amino acid substitutions responsible for either decreasing or increasing the stability of the related protein via cross validation methods on a large dataset of single amino acid mutations. No tertiary structure of the protein is required for the prediction. (Cheng et al., 2006). MUpro web server available at (http://mupro.proteomics.ics.uci.edu/) was used in this study and a confidence score <0 was predicted to cause decrease in the stability and a score >0 is predicted to cause increase in the stability.

### 2.5. Structural conformation and conservation analysis by ConSurf sever

Highly conserved functional regions of the protein coded by CDKN1A gene was identified by ConSurf tool (http://consurf.tau.ac.il/). It constructs a phylogenetic tree, following multiple sequence alignment of a query sequence, to run a high-throughput identification process of the functional regions of a protein based on evolutionary data (Ashkenazy et al., 2016).

### 2.6. Prediction of secondary structure by PSIPRED

Secondary structure of CDKN1A was predicted using PSIPRED server available at (http://bioinf.cs.ucl.ac.uk/psipred/) (Buchan et al., 2013). It is based on a two stage neural network with the implementation of position specific scoring matrices constructed from PSI-BLAST to predict the available secondary structures of a protein. (Jones, 1999).

### 2.7. Homology Modelling

3D structures of the protein encoded by CDKN1A was constructed using four different homology modelling tools as no crystal structure with appropriate length of this protein was available in protein data bank.

#### 2.7.1. Homology modelling by Muster server

MUlti-score ThreadER (MUSTER) algorithm (https://zhanglab.ccmb.med.umich.edu/MUSTER/), with the implementation of MODELLER v8.2 to construct full length protein model following sequence-template alignment, was used in this study. It uses information such as-sequence profiles, secondary structures, structure fragment profiles, solvent accessibility, dihedral torsion angles, hydrophobic scoring matrix of different sequences to construct the model (Wu and Zhang, 2008). It provides several template based models with a scoring function expressed as Z value to determine the best model.

#### 2.7.2. Homology modelling by Phyre-2 server

Phyre-2 tool (http://www.sbg.bio.ic.ac.uk/phyre2/), based on Hidden Markov method, was used to predict the homology based three dimensional structure of the query amino acid sequence. It combines four steps to build a model: (1) collecting homologous sequence, (2) screening of fold library, (3) loop modelling, (4) multiple template modelling by ab-initio folding simulation-Poing and (4) Side chain placement (Kelley et al., 2015).

#### 2.7.3. Homology modelling by RaptorX server

RaptorX server available at (http://raptorx.uchicago.edu/StructurePrediction/predict/), using RaptorX-Boost and RaptorX-MSA to construct three dimensional structure of a protein, predicted the 3D model of the protein coded by CDKN1A. It combines a nonlinear scoring function and a probabilistic-consistency algorithm for predicting the model structure (Källberg et al., 2012).

#### 2.7.4. Homology modelling by Swiss-Model server

Swiss-Model workplace available at (https://swissmodel.expasy.org/), a web based tool for the homology modelling of protein, was used to avail the three dimensional structure of CDKN1A (Arnold et al., 2006). The accuracy of the constructed model is calculated by CAMEO system. Swiss-model is an automated tool based on evolutionary information which searches for the best sequence-template alignment from its high-throughput template library (SMTL) to build the model (Biasini et al., 2014).

### 2.8. Model Refinement, energy minimization and mutation

High resolution structure refinement of the protein model at the atomic level was carried out by ModRefiner tool (Xu and Zhang, 2011) and energy minimization executed for the improvement of the quality of models with the GROMOS 96 forcefield (van Gunsteren, 1996) implementation of DeepView v.4.1.0 (swiss pdb viewer) (Guex and Peitsch, 1997). Execution of energy minimization was done in vacao without any reaction field. Swiss PdbViewer v.4.1.0 was also implemented to build mutated models by browsing a rotamer library of the protein model. Energy minimization of mutant models was brought about to increase the quality.

### 2.9. Visualization of different models by BIOVIA Discovery Studio Visualizer 2016

Discovery Studio Visualizer (*Dassault Systèmes*, 2016) tool was used to display the 3D structures of the constructed models following structure refinement and energy minimization.

### 2.10. Validation of models

#### 2.10.1. Ramachandran plot analysis

Ramachandran plot analysis of the protein models, available at ((http://www-cryst.bioc.cam.ac.uk/rampage) determined the energetically allowed sites of amino acid residues in a protein structure through the calculation of backbone dihedral angles of ψ against φ of amino acid residues (Lovell et al., 2003).

#### 2.10.2. QMEAN6 Z-value

The degree of nativeness of the predicted models are evaluated by the QMEAN scoring (https://swissmodel.expasy.org/qmean/). The major five geometrical features of the protein is predicted by this composite scoring function (Benkert et al., 2008).

### 2.11. Screening of most frequent deleterious substitution

IMB SPSS Statistics for Windows v.20 (*IBM Corp*., 2011) was used to find the most frequent variants having deleterious effect based on the prediction of six evaluation methods-Predict SNP, MAPP, PhD SNP, SNAP, I-Mutant 3.0 and Mupro, following the initial screening by SIFT and Polyphen-2, which predicted 53 nsSNPs as being harmful out of 118.

### 2.12. Visualization of selected mutations using Mutation 3D server

Mutation 3D server (http://mutation3d.org/), an interactive tool to visualize amino acid substitutions along with their position on functional domains, was employed in this study. The algorithm used here is an approach based on 3D clustering method and was used to predict driver genes in cancer (Meyer et al., 2016).

### 2.13. Prediction of biophysical activity by Align-GVGD

Align-GVGD predicted the biophysical characteristics of the amino acid substitutions (http://agvgd.hci.utah.edu/). Grantham Variation (GV) and Grantham Deviation (GD) scores (0 to >200) and graded classifiers (C0 to C65) were calculated to predict either deleterious or neutral variants (Mathe et al., 2006). An extension of Grantham difference to multiple sequence alignments and true simultaneous multiple comparisons, Align-GVGD, predicts missense substitutions (Tavtigian et al., 2006).

### 2.14. Harmful effects of selected variants by MutPred2 server

MutPred2 (http://mutpred2.mutdb.org/) web server proposed the reasons behind the disease related amino acid substitution at the molecular level (Pejaver et al., 2017). The result generated by this algorithm predicts a general probability score to define a substitution as deleterious or disease related and includes top five molecular features with a P-value. Scores are interpreted in three categories-actionable hypotheses (general probability score, *g* > 0.5 and property score, *p* < 0.05), confident hypotheses (*g* > 0.75 and *p* < 0.05), and very confident hypotheses (*g* > 0.75 and *p* < 0.01) (Li et al., 2009).

## 3. Results

NCBI dbSNP database contains information on SNPs of various genes. Mining this database on December 11, 2017, revealed 1766 SNPs of CDKN1A gene, of which 1718 were from *Homo sapiens*. Of the 118 CDKN1A nonsynonymous SNPs found in humans, 113 were missense mutation and the rest 5 were nonsense mutation. FASTA format of the protein encoded by CDKN1A gene, retrieved from Ensemble database, is 164 amino acid long.

### 3.1. Prediction of functional context of nsSNPs

Functional features of the 118 CDKN1A nsSNPs analyzed by SIFT and Polyphen-2 screened out 65 nsSNPs as because they were predicted as “Tolerated” by SIFT and “Benign” by Polyphen-2 server. All 5 nonsense mutations were found to be either tolerated or benign by these servers.

**Table 1** represents the final scores of 53 nsSNPs generated by the sequence homology based software SIFT and Polyphen-2. In SIFT, 23 nsSNPs out of 53 had a tolerance index ≤0.05 and predicted as “Damaging”. While analyzing with Polyphen-2, two score were calculated-HumDiv and HumVar. Polyphen-2 gives score ranging from 0 (neutral) to 1 (damaging). HumDiv predicted 34 nsSNPs as “Probably Damaging (High Confidence)” and 12 as “Possibly Damaging (Low confidence)” out of 53 nsSNPs. HumVar reported 28 nsSNPs as “Probably Damaging (High Confidence)” and 10 as “Possibly Damaging (Low confidence)” out of 53 nsSNPs.

**Table 1.**
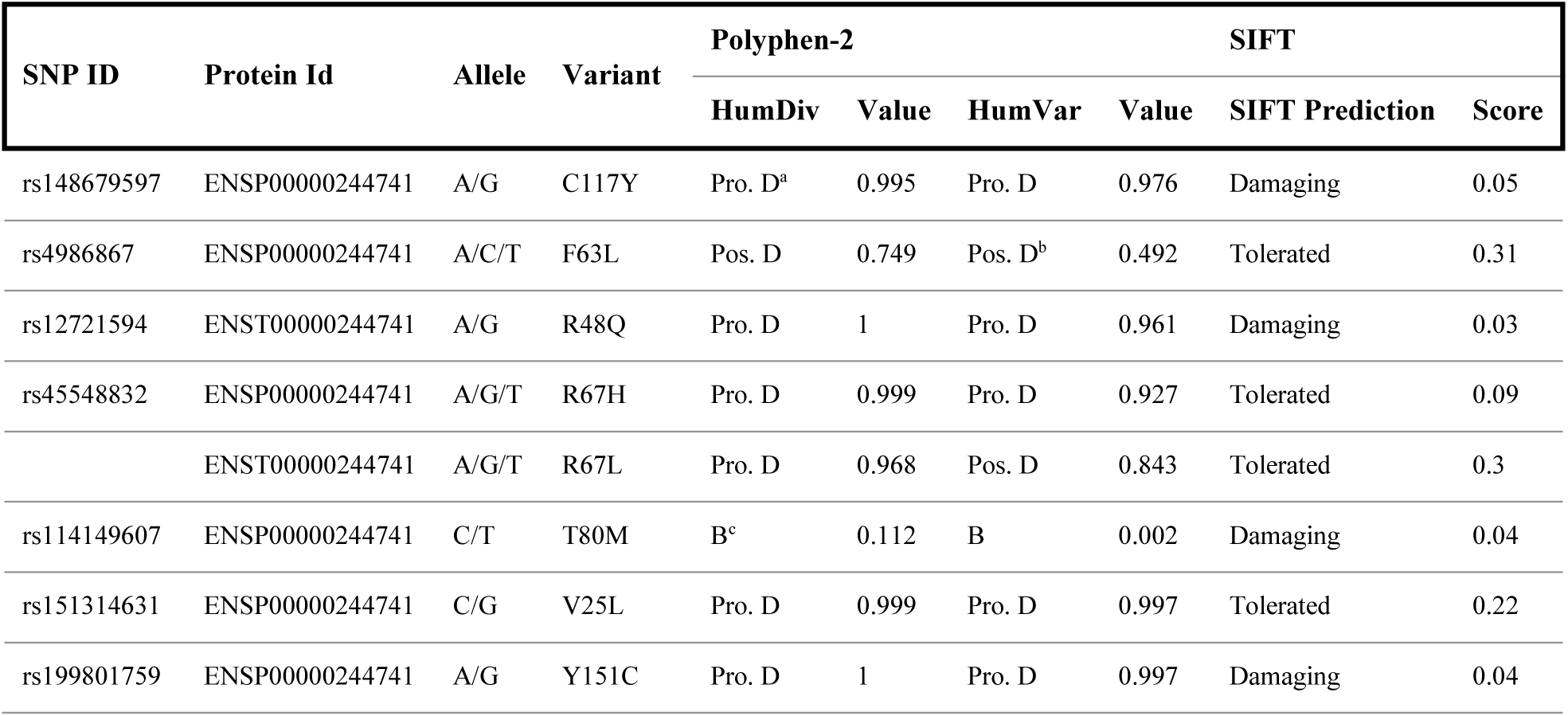
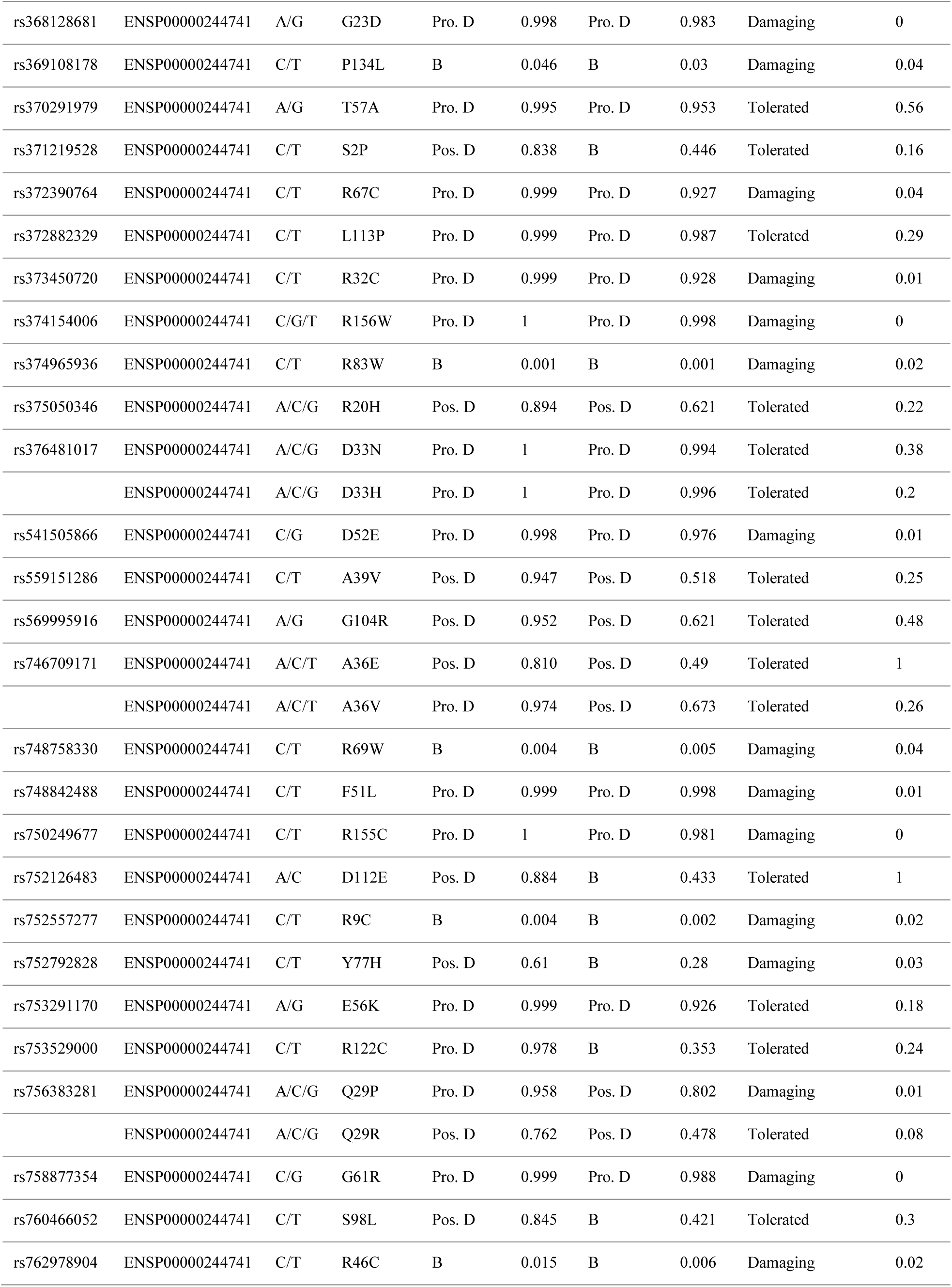
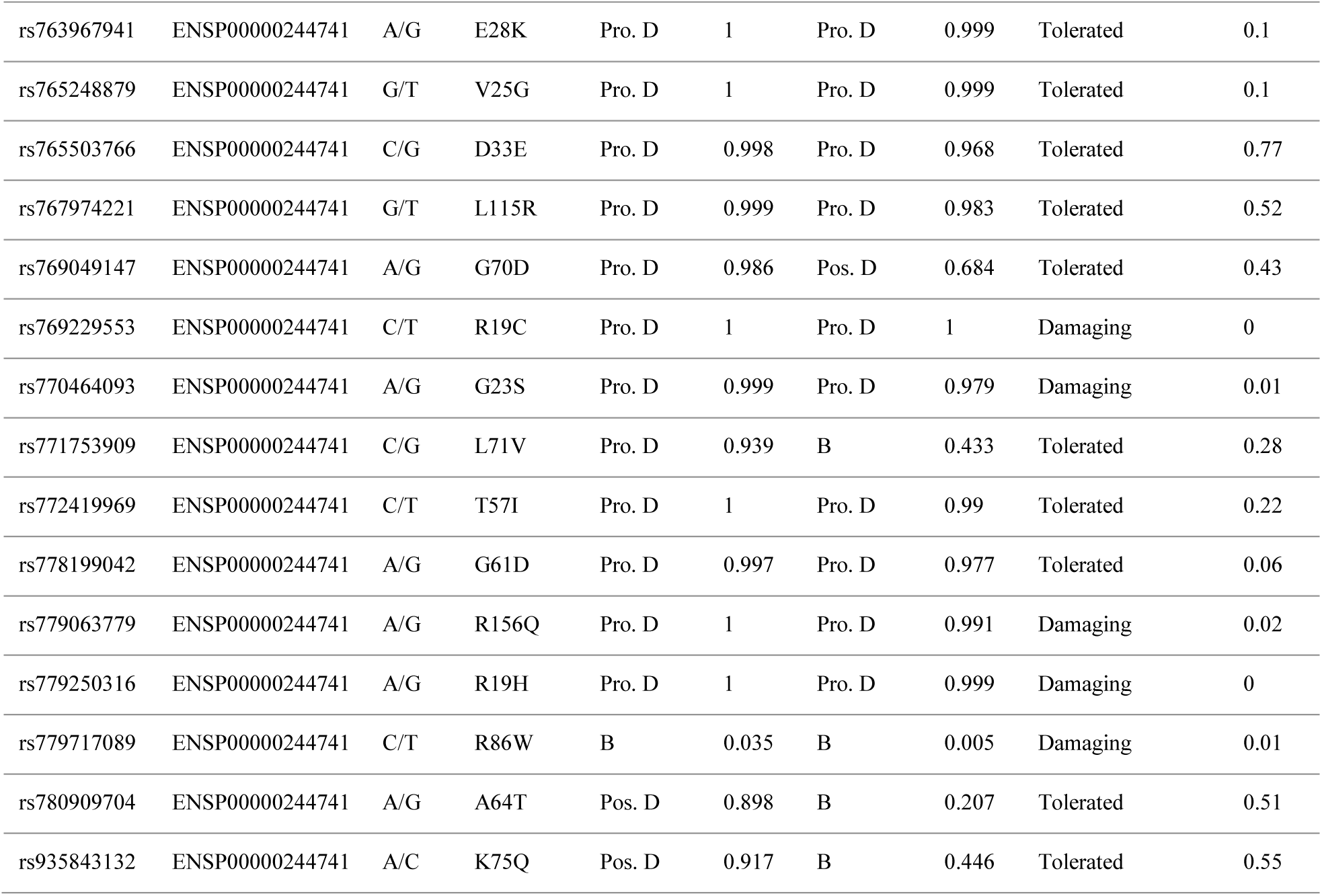
Screening of Deleterious Single Nucleotide Polymorphism (SNP) Predicted by SIFT and Polyphen-2. ^a^ Probably damaging (more confident); ^b^ Possibly damaging (less confident); ^c^ Benign.

| SNP ID | Protein Id | Allele | Variant | Polyphen-2 |  |  | SIFT |  |  |
| --- | --- | --- | --- | --- | --- | --- | --- | --- | --- |
|  |  |  |  | HumDiv | Value | HumVar | Value | SIFT Prediction | Score |
| rs148679597 | ENSP00000244741 | A/G | C117Y | Pro. D<sup>a</sup> | 0.995 | Pro. D | 0.976 | Damaging | 0.05 |
| rs4986867 | ENSP00000244741 | A/C/T | F63L | Pos. D | 0.749 | Pos. D<sup>b</sup> | 0.492 | Tolerated | 0.31 |
| rs12721594 | ENST00000244741 | A/G | R48Q | Pro. D | 1 | Pro. D | 0.961 | Damaging | 0.03 |
| rs45548832 | ENSP00000244741 | A/G/T | R67H | Pro. D | 0.999 | Pro. D | 0.927 | Tolerated | 0.09 |
|  | ENST00000244741 | A/G/T | R67L | Pro. D | 0.968 | Pos. D | 0.843 | Tolerated | 0.3 |
| rs114149607 | ENSP00000244741 | C/T | T80M | B<sup>c</sup> | 0.112 | B | 0.002 | Damaging | 0.04 |
| rs151314631 | ENSP00000244741 | C/G | V25L | Pro. D | 0.999 | Pro. D | 0.997 | Tolerated | 0.22 |
| rs199801759 | ENSP00000244741 | A/G | Y151C | Pro. D | 1 | Pro. D | 0.997 | Damaging | 0.04 |
| rs368128681 | ENSP00000244741 | A/G | G23D | Pro. D | 0.998 | Pro. D | 0.983 | Damaging | 0 |
| rs369108178 | ENSP00000244741 | C/T | P134L | B | 0.046 | B | 0.03 | Damaging | 0.04 |
| rs370291979 | ENSP00000244741 | A/G | T57A | Pro. D | 0.995 | Pro. D | 0.953 | Tolerated | 0.56 |
| rs371219528 | ENSP00000244741 | C/T | S2P | Pos. D | 0.838 | B | 0.446 | Tolerated | 0.16 |
| rs372390764 | ENSP00000244741 | C/T | R67C | Pro. D | 0.999 | Pro. D | 0.927 | Damaging | 0.04 |
| rs372882329 | ENSP00000244741 | C/T | L113P | Pro. D | 0.999 | Pro. D | 0.987 | Tolerated | 0.29 |
| rs373450720 | ENSP00000244741 | C/T | R32C | Pro. D | 0.999 | Pro. D | 0.928 | Damaging | 0.01 |
| rs374154006 | ENSP00000244741 | C/G/T | R156W | Pro. D | 1 | Pro. D | 0.998 | Damaging | 0 |
| rs374965936 | ENSP00000244741 | C/T | R83W | B | 0.001 | B | 0.001 | Damaging | 0.02 |
| rs375050346 | ENSP00000244741 | A/C/G | R20H | Pos. D | 0.894 | Pos. D | 0.621 | Tolerated | 0.22 |
| rs376481017 | ENSP00000244741 | A/C/G | D33N | Pro. D | 1 | Pro. D | 0.994 | Tolerated | 0.38 |
|  | ENSP00000244741 | A/C/G | D33H | Pro. D | 1 | Pro. D | 0.996 | Tolerated | 0.2 |
| rs541505866 | ENSP00000244741 | C/G | D52E | Pro. D | 0.998 | Pro. D | 0.976 | Damaging | 0.01 |
| rs559151286 | ENSP00000244741 | C/T | A39V | Pos. D | 0.947 | Pos. D | 0.518 | Tolerated | 0.25 |
| rs569995916 | ENSP00000244741 | A/G | G104R | Pos. D | 0.952 | Pos. D | 0.621 | Tolerated | 0.48 |
| rs746709171 | ENSP00000244741 | A/C/T | A36E | Pos. D | 0.810 | Pos. D | 0.49 | Tolerated | 1 |
|  | ENSP00000244741 | A/C/T | A36V | Pro. D | 0.974 | Pos. D | 0.673 | Tolerated | 0.26 |
| rs748758330 | ENSP00000244741 | C/T | R69W | B | 0.004 | B | 0.005 | Damaging | 0.04 |
| rs748842488 | ENSP00000244741 | C/T | F51L | Pro. D | 0.999 | Pro. D | 0.998 | Damaging | 0.01 |
| rs750249677 | ENSP00000244741 | C/T | R155C | Pro. D | 1 | Pro. D | 0.981 | Damaging | 0 |
| rs752126483 | ENSP00000244741 | A/C | D112E | Pos. D | 0.884 | B | 0.433 | Tolerated | 1 |
| rs752557277 | ENSP00000244741 | C/T | R9C | B | 0.004 | B | 0.002 | Damaging | 0.02 |
| rs752792828 | ENSP00000244741 | C/T | Y77H | Pos. D | 0.61 | B | 0.28 | Damaging | 0.03 |
| rs753291170 | ENSP00000244741 | A/G | E56K | Pro. D | 0.999 | Pro. D | 0.926 | Tolerated | 0.18 |
| rs753529000 | ENSP00000244741 | C/T | R122C | Pro. D | 0.978 | B | 0.353 | Tolerated | 0.24 |
| rs756383281 | ENSP00000244741 | A/C/G | Q29P | Pro. D | 0.958 | Pos. D | 0.802 | Damaging | 0.01 |
|  | ENSP00000244741 | A/C/G | Q29R | Pos. D | 0.762 | Pos. D | 0.478 | Tolerated | 0.08 |
| rs758877354 | ENSP00000244741 | C/G | G61R | Pro. D | 0.999 | Pro. D | 0.988 | Damaging | 0 |
| rs760466052 | ENSP00000244741 | C/T | S98L | Pos. D | 0.845 | B | 0.421 | Tolerated | 0.3 |
| rs762978904 | ENSP00000244741 | C/T | R46C | B | 0.015 | B | 0.006 | Damaging | 0.02 |
| rs763967941 | ENSP00000244741 | A/G | E28K | Pro. D | 1 | Pro. D | 0.999 | Tolerated | 0.1 |
| rs765248879 | ENSP00000244741 | G/T | V25G | Pro. D | 1 | Pro. D | 0.999 | Tolerated | 0.1 |
| rs765503766 | ENSP00000244741 | C/G | D33E | Pro. D | 0.998 | Pro. D | 0.968 | Tolerated | 0.77 |
| rs767974221 | ENSP00000244741 | G/T | L115R | Pro. D | 0.999 | Pro. D | 0.983 | Tolerated | 0.52 |
| rs769049147 | ENSP00000244741 | A/G | G70D | Pro. D | 0.986 | Pos. D | 0.684 | Tolerated | 0.43 |
| rs769229553 | ENSP00000244741 | C/T | R19C | Pro. D | 1 | Pro. D | 1 | Damaging | 0 |
| rs770464093 | ENSP00000244741 | A/G | G23S | Pro. D | 0.999 | Pro. D | 0.979 | Damaging | 0.01 |
| rs771753909 | ENSP00000244741 | C/G | L71V | Pro. D | 0.939 | B | 0.433 | Tolerated | 0.28 |
| rs772419969 | ENSP00000244741 | C/T | T57I | Pro. D | 1 | Pro. D | 0.99 | Tolerated | 0.22 |
| rs778199042 | ENSP00000244741 | A/G | G61D | Pro. D | 0.997 | Pro. D | 0.977 | Tolerated | 0.06 |
| rs779063779 | ENSP00000244741 | A/G | R156Q | Pro. D | 1 | Pro. D | 0.991 | Damaging | 0.02 |
| rs779250316 | ENSP00000244741 | A/G | R19H | Pro. D | 1 | Pro. D | 0.999 | Damaging | 0 |
| rs779717089 | ENSP00000244741 | C/T | R86W | B | 0.035 | B | 0.005 | Damaging | 0.01 |
| rs780909704 | ENSP00000244741 | A/G | A64T | Pos. D | 0.898 | B | 0.207 | Tolerated | 0.51 |
| rs935843132 | ENSP00000244741 | A/C | K75Q | Pos. D | 0.917 | B | 0.446 | Tolerated | 0.55 |

### 3.2. Prediction of deleterious and disease related amino acid substitutions

Predict-SNP is a consensus classifiers for the prediction of disease related mutations. Analysis of 53 of nsSNPs by this server also predicted MAPP, PhD-SNP and SNAP scores of nsSNPs. All results are summarized in

**Table *2***. Out of this 53 nsSNPs, 31, 50, 19 and 29 were predicted as “Deleterious” by Predict-SNP, MAPP, PhD-SNP and SNAP respectively.

**Table 2.**
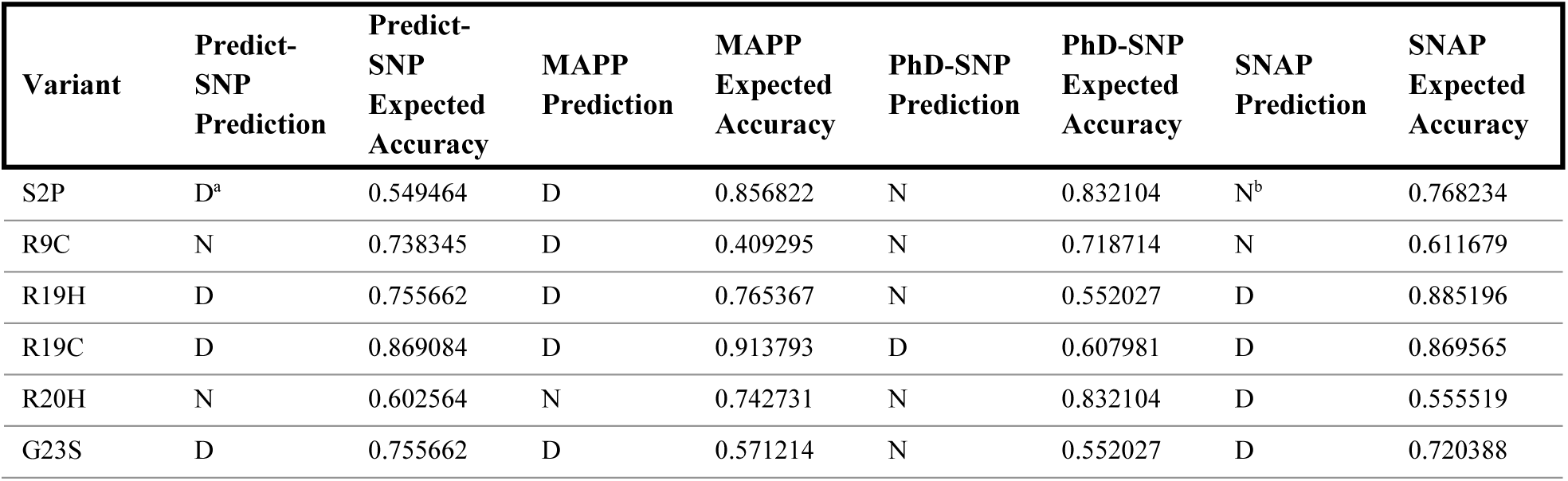
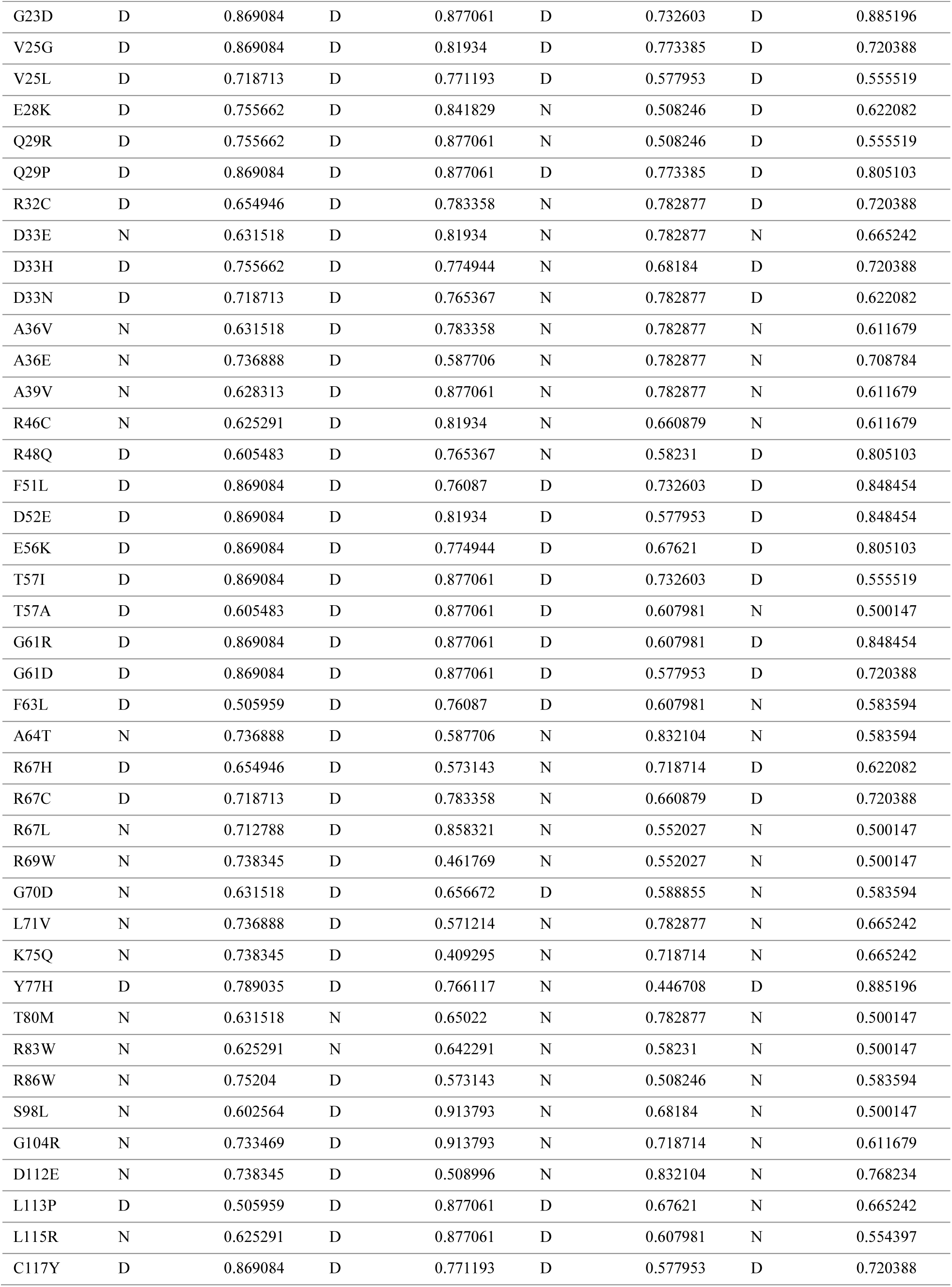
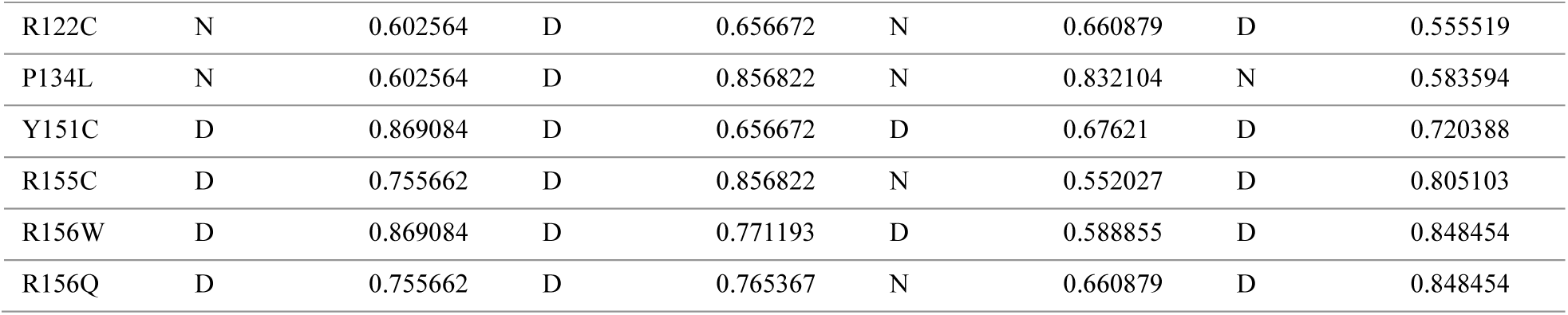
Prediction of Disease Related Mutations by Predict-SNP, MAPP, PhD-SNP, and SNAP. ^a^ Deleterious effect; ^b^ Neutral effect.

### 3.3. I-mutant 3.0 and Mupro analysis of protein’s stability change upon amino acid substitution

Prediction of the changes in protein’s stability upon amino acid substitution performed by I-Mutant 3.0 server and Mupro Server predicted 46 and 41 amino acid substitution, out of 53, respectively, to bring decrease in the stability of the cyclin dependent kinase 1 protein encoded by CDKN1A gene. All Scores from these servers are represented in

**Table 3**. Stability change in I-Mutant 3.0 was calculated at 25 ºC and pH 7 and expressed as DDG value=DG (new protein)-DG (wild type) in kcal/mol.

**Table 3.**
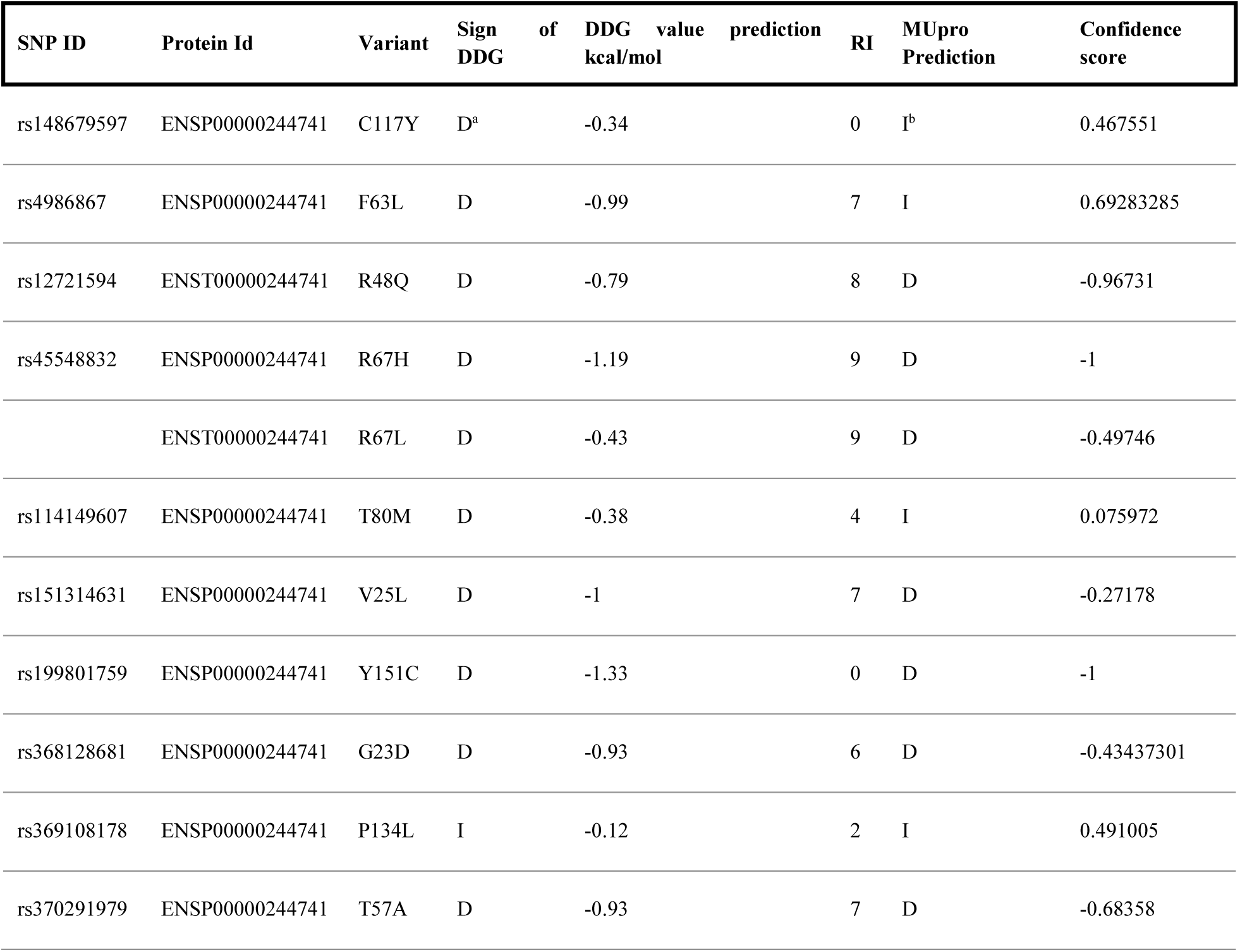
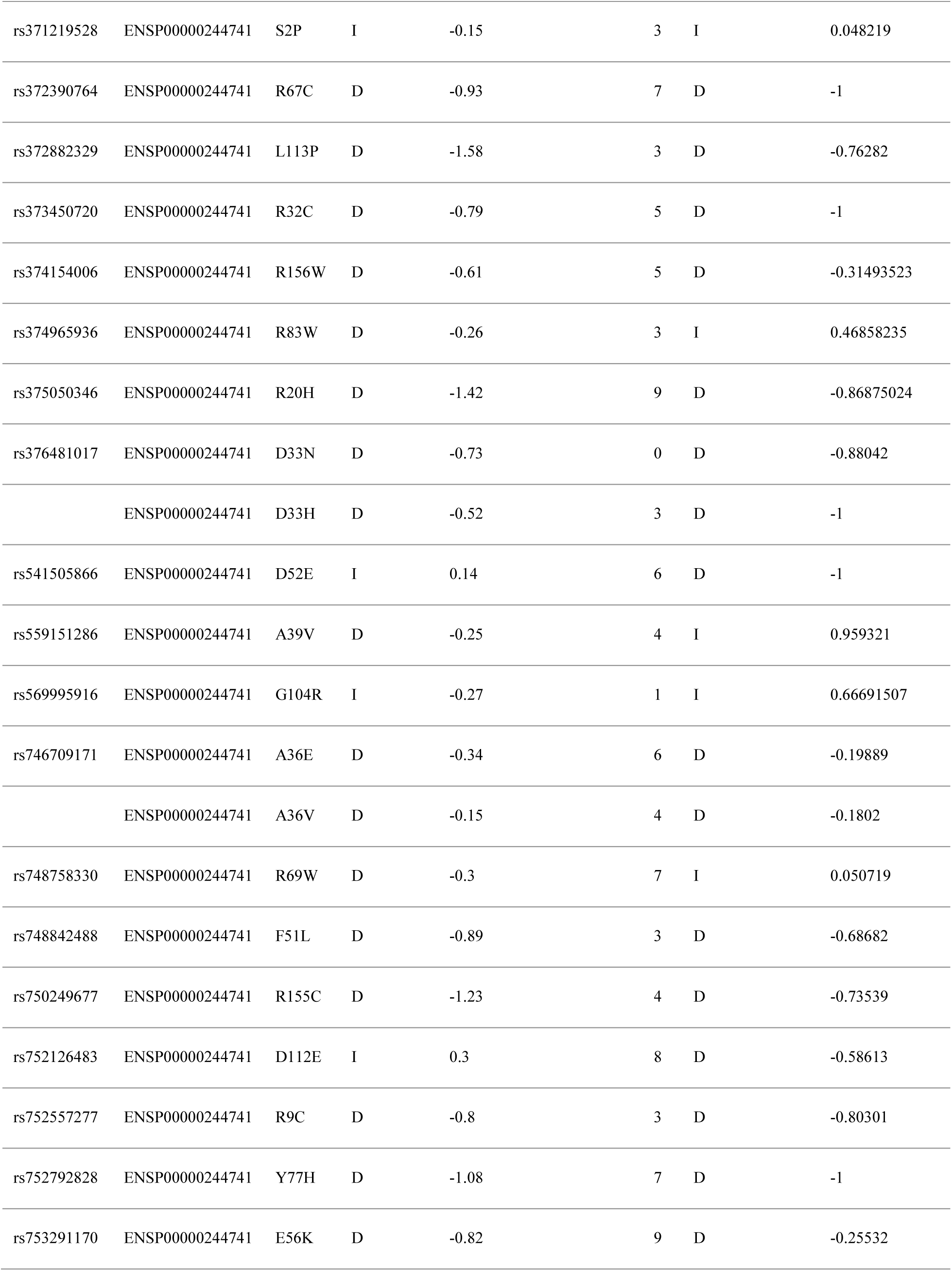
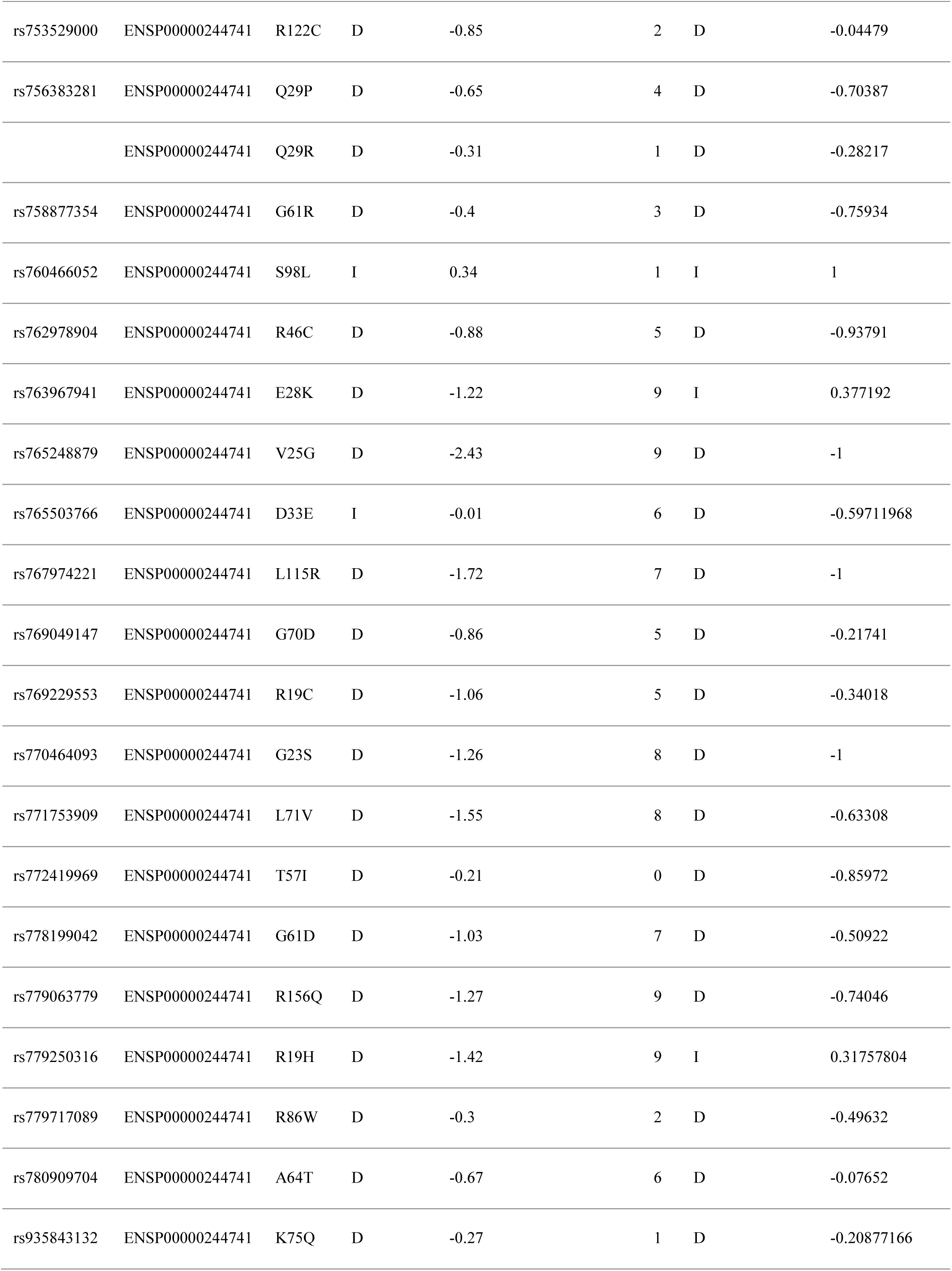
Characterization of the Effect of nsSNPs on Protein Stability by I-Mutant 3.0 and MUpro. ^a^ Decrease the stability of protein structure; ^b^ Increase the stability of protein structure.

### 3.4. Screening of most frequent deleterious amino acid substitution

Following the screening of harmful nsSNPs by SIFT and Polyphen-2 server, the rest 53 nsSNPs, predicted to be deleterious, were further analyzed. A distribution of harmful substitution predicted by Predict SNP, MAPP, PhD-SNP, SNAP, I-Mutant 3.0 and Mupro is shown in **Figure 1**. Disease related mutations exposed by Predict SNP, MAPP, PhD-SNP and SNAP predicted 58.5%, 93.3%, 35.8%, and 54.7% nsSNPs, respectively, as deleterious. Analysis by I-Mutant 3.0 and Mupro suggested that 86.8% and 77.4% nsSNPs, respectively, were affected by amino acid substitution and resulted in decrease in stability. Frequency of deleterious nsSNPs in various servers was calculated by IBM SPSS v20 and Microsoft office 2013.

**Figure 1.**
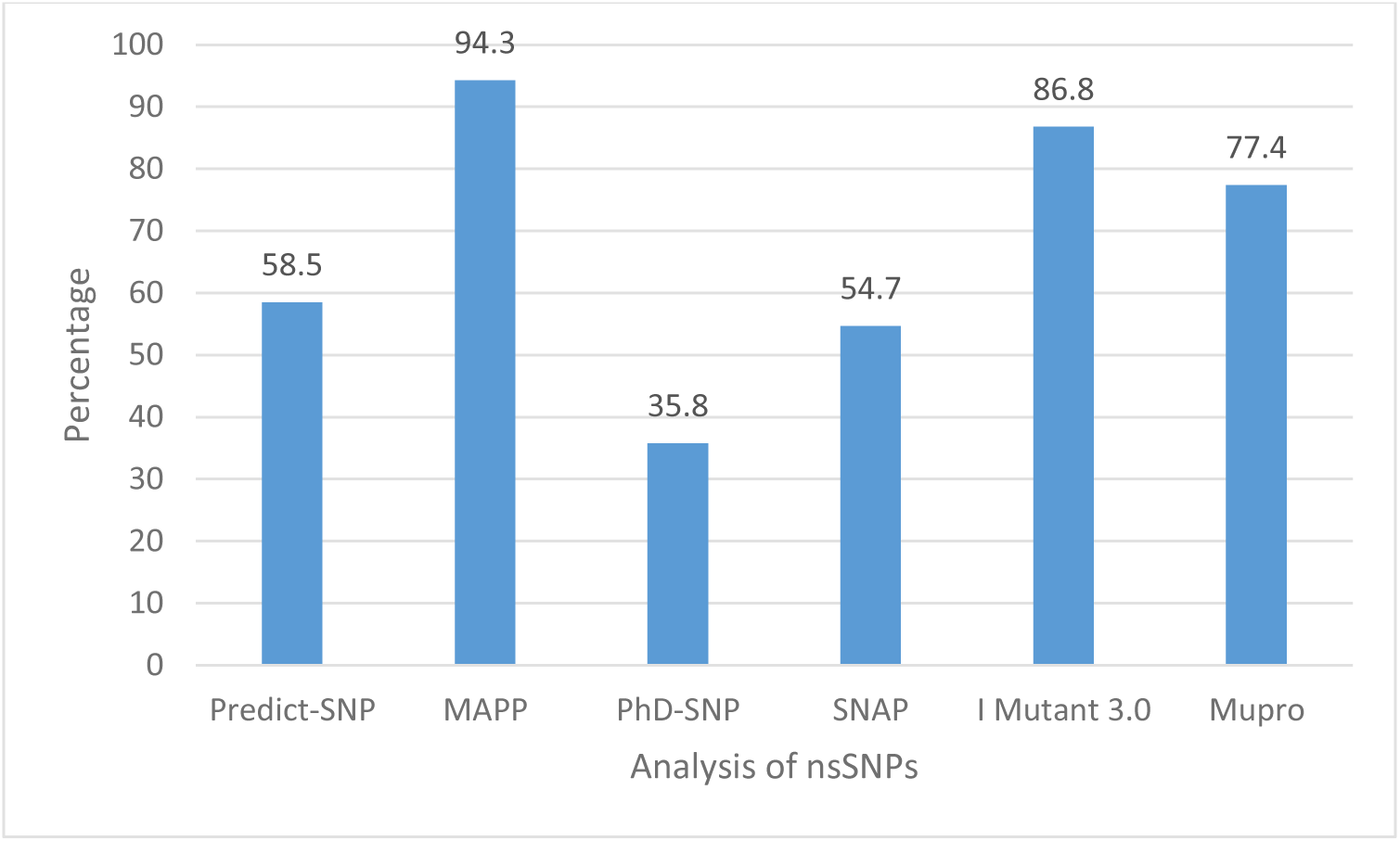
Overview of the analysis of deleterious and stability decreased cyclin dependent kinase inhibitor 1 due to nsSNPs by various in silico tools

Prediction and distribution of deleterious nsSNPs by all six methods-Predict-SNP, MAPP, PhD-SNP, SNAP, I-Mutant 3.0 and Mupro, illustrated in **Figure 2**, revealed the most frequent 12 amino acid substitution predicted to be either “Deleterious” or “Stability Decreasing”. Amino acid substitution-R19C, G23D, V25G, V25L, Q29P, F51L, E56K, T57I, G61R, G61D, Y151C and R156W were predicted as most harmful by all the above methods.

**Figure 2.**
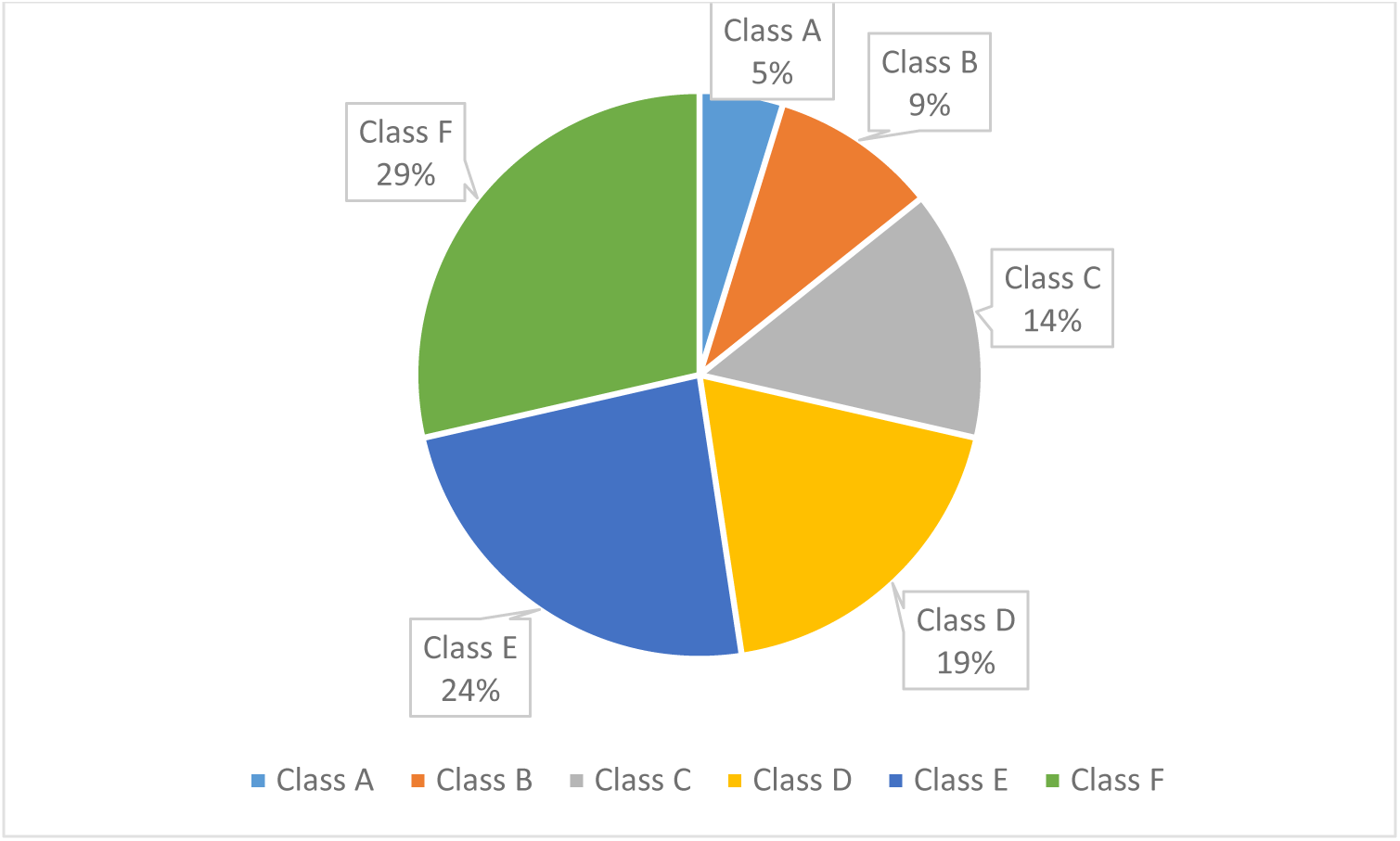
Prediction and distribution of deleterious nsSNPs by Predict-SNP, MAPP, PhD-SNP, SNAP, I-Mutant 3.0 and Mupro methods; Class A, predicted damaging by at least one method, T80M, R83W, S98L, G104R and P134L; Class B, predicted damaging by at least two methods, D33E,A39V, R69W, D112E; Class C, predicted damaging by at least three methods, R9C, R20H, A36V, A36E, R46C, A64T, R67L, L71V, R86W; Class D, predicted damaging by at least four methods, S2P, R19H, E28K, F63L, G70D, K75Q, L115R, R122C; Class E, predicted damaging by at least five methods, G23S, Q29R, R32C, D33H, D33N, R48Q, D52E, T57A, R67H, R67C, Y77H, L113P, C117Y, R155C, R156Q; Class F, predicted damaging by all six methods, R19C, G23D, V25G, V25L, Q29P, F51L, E56K, T57I, G61R, G61D, Y151C, R156W.

### 3.5. Structural conformation and conservation analysis by ConSurf sever

A colorimetric conservation score was produced as result by the ConSurf server (**Figure 3**). Highly conserved functional region of the protein was revealed by ConSurf tool. It was found that R19C, G23D, F51L, Y151C and R156W have a conservation score of 9; V25G, V25L, Q29P, T57I, G61R and G61D have a conservation score of 8; E56K have a conservation score of 7.

**Figure 3.**
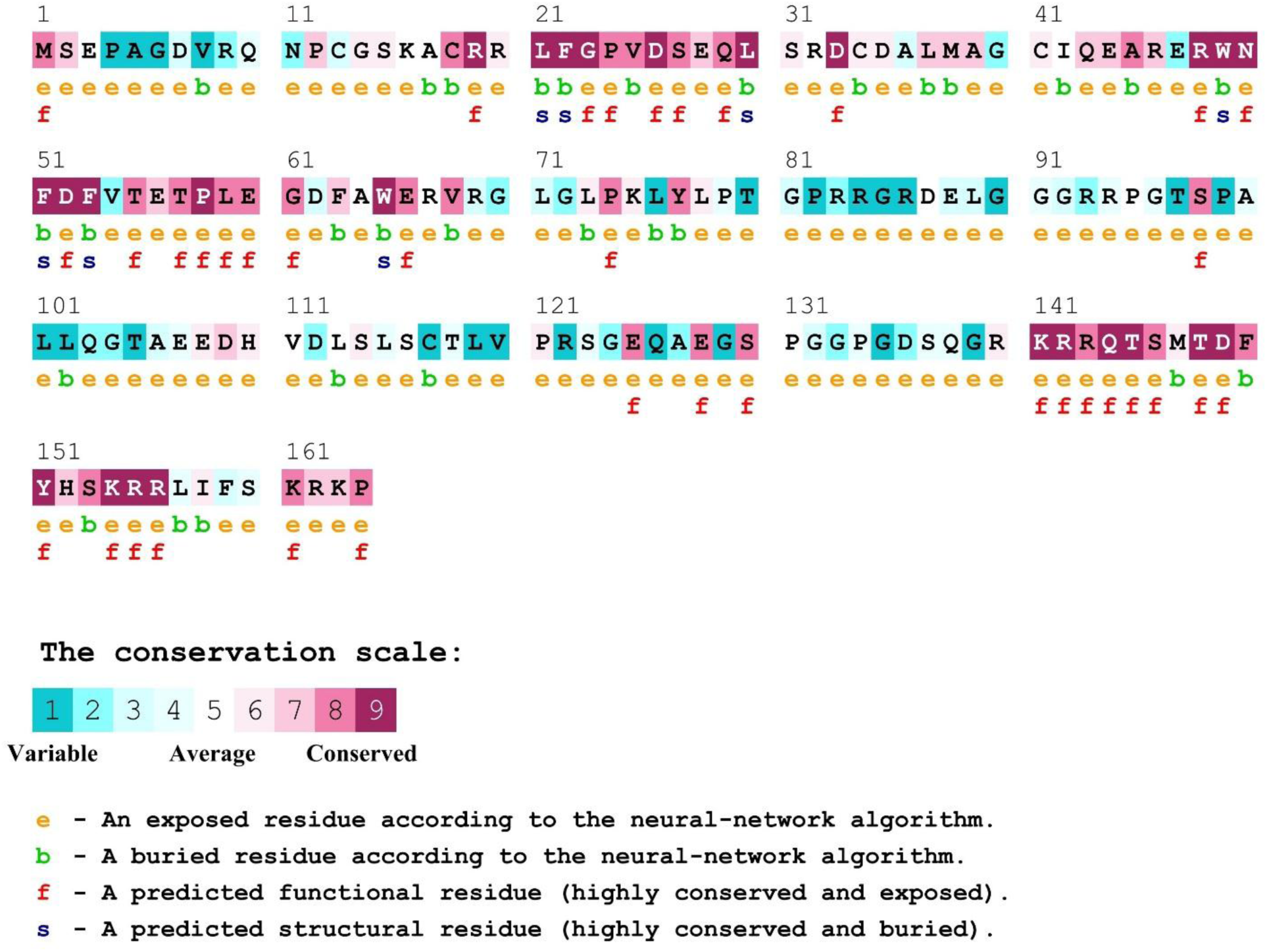
Prediction of evolutionary conserved amino acid residues by ConSurf server. Conservation score is represented as the color coding bars.

### 3.6. Prediction of secondary structure by PSIPRED

Distribution of alpha helix and beta sheet and coil is exposed by PSIPRED (**Figure 4**). Among the secondary structures, the highest in percentage was coils (82.93%) followed by alpha helix (14.02%) and beta sheet (3.05%).

**Figure 4.**
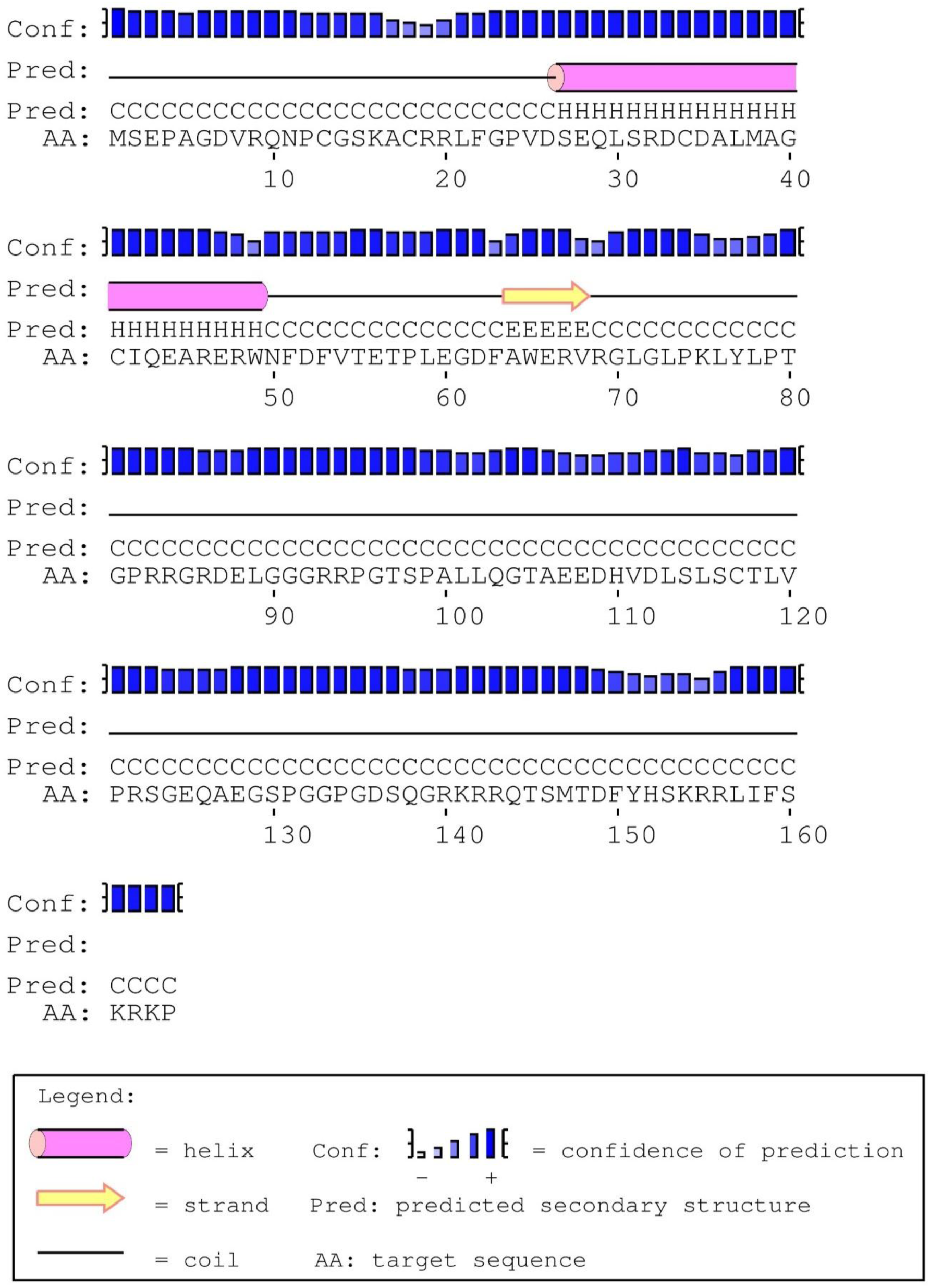
Prediction of secondary structure of by PSIPRED server

### 3.7. Homology Modelling

Structure refined and energy minimized homology models of CDKN1A by MUSTER, Phyre-2, RaptorX and Swiss model serves are illustrated using computational program Discovery Studio Modeling Environment 2016 (**Figure 5**). These models were checked for validation through Ramachandran plot analysis and QMEAN Z scores.

**Figure 5.**
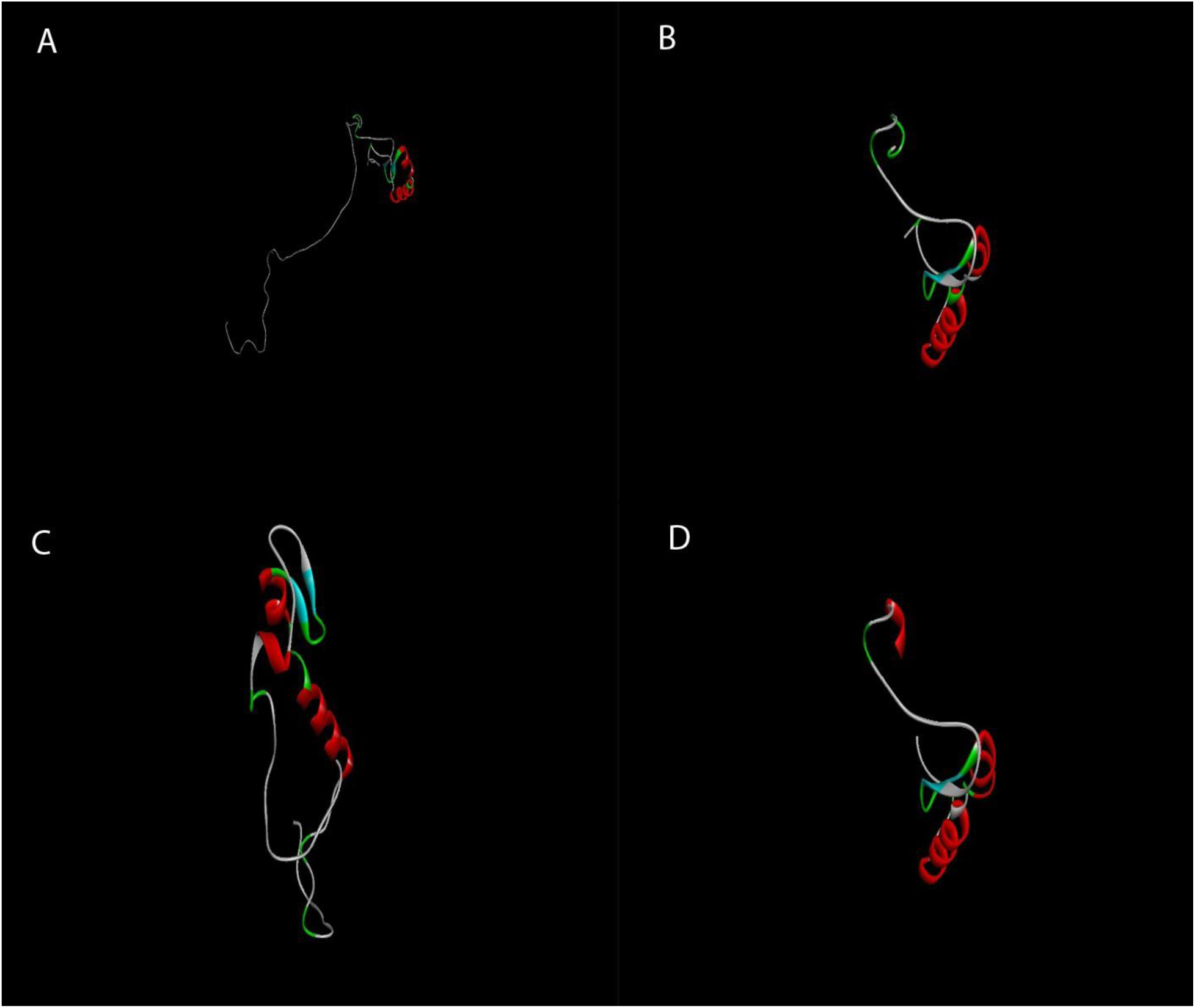
Homology models from different servers; A. CDKN1A_Muster, B. CDKN1A_Phyre2, C. CDKN1A_RaptorX, D. CDKN1A_Swiss model

#### 3.7.1. Ramachandran plot analysis

Ramachandran plot is an x-y plot of phi/psi dihedral angles between NC-alpha and Calpha-C bonds to evaluate a protein’s backbone conformation. Ramachandran plots of structure refined and energy minimized homology models are illustrated in **Figure 6**.

**Figure 6.**
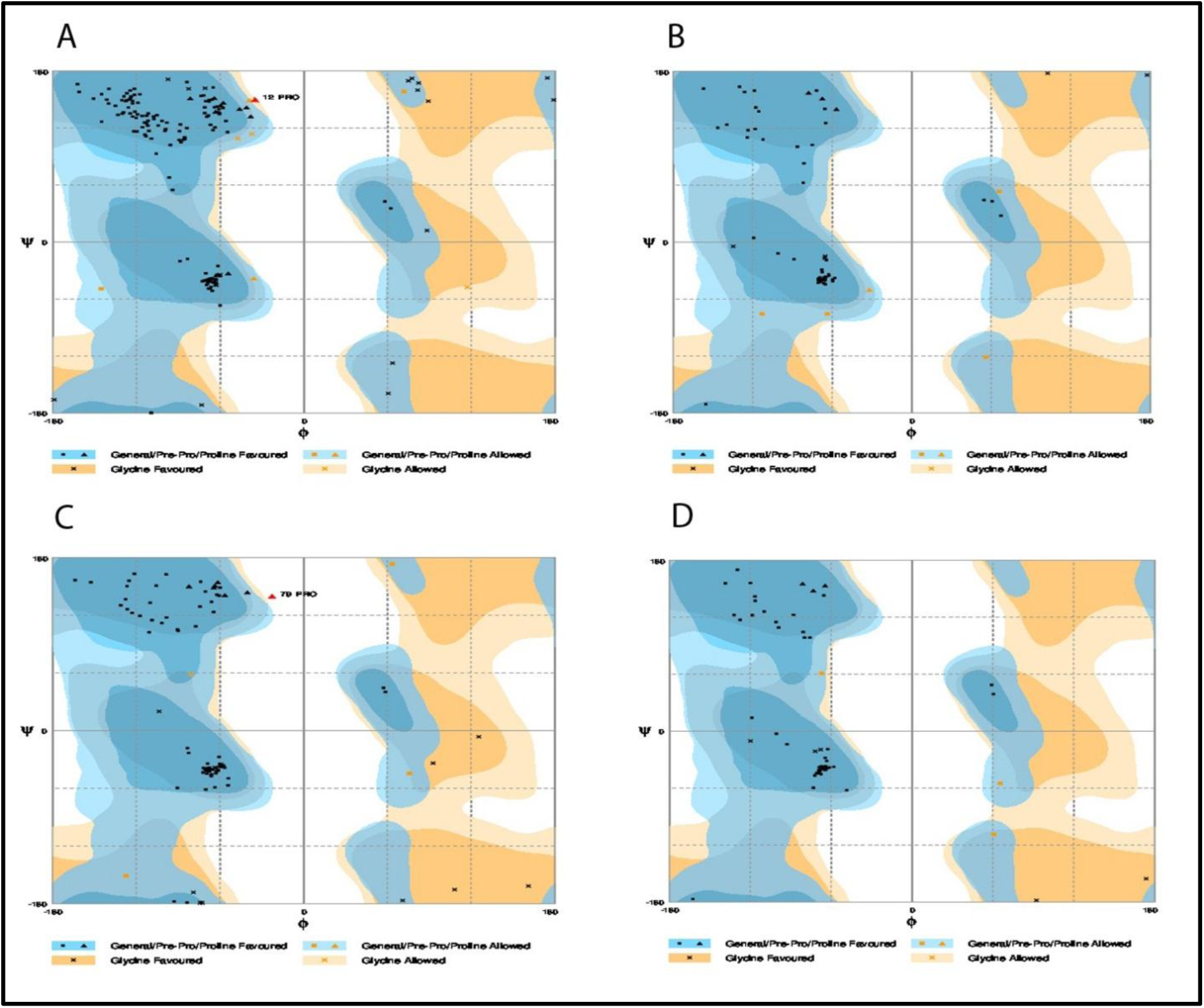
Ramachandran plots of different models; A. CDKN1A_Muster, B. CDKN1A_Phyre2, C. CDKN1A_RaptorX, D. CDKN1A_Swiss model

**Table 4** represents the number of residues of different models (structure refined and energy minimized) in favored, allowed and outlier regions generated by the Ramachandran plot. Model generated by MUSTER had highest number of residues (95.1%) in favored region followed by Swiss model (95%), RaptorX (94.5%) and Phyre-2 (92.3%) server.

**Table 4.** Ramachandran plot analysis of different homology models of CDKN1A

| Model | Residues in favored region |  | Residues in allowed region |  | Residues in outlier region |  |
| --- | --- | --- | --- | --- | --- | --- |
|  | No of residues | % of residues | No of residues | % of residues | No of residues | % of residues |
| CDKN1A_muster | 154 | 95.1 | 7 | 4.3 | 1 | 0.6 |
| CDKN1A_phyre 2 | 60 | 92.3 | 5 | 7.7 | 0 | 0 |
| CDKN1A_raptor X | 86 | 94.5 | 4 | 4.4 | 1 | 1.1 |
| CDKN1A_swiss model | 57 | 95.0 | 3 | 5.0 | 0 | 0 |

#### 3.7.2. QMEAN6 Z-value

**Table 5** shows the Z scores obtained from QMEAN Server (Swiss Model) using structure refined and energy minimized models. Overall, the total QMEAN score (Z value) of the models were better for RaptorX (-1.68), followed by Swiss model (-2.01), Phyre-2 (-2.12) and MUSTER (-2.30).

**Table 5.**
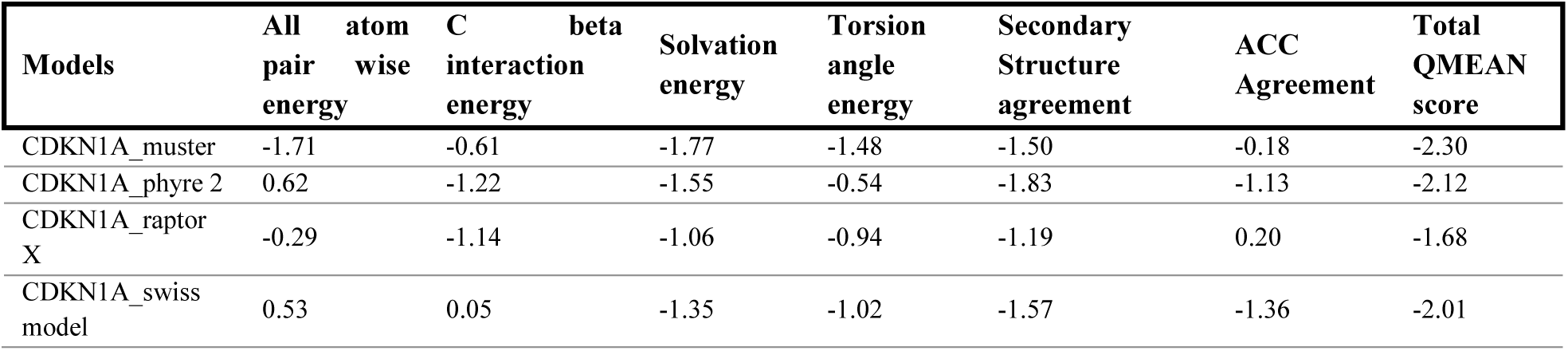
Z scores obtained from QMEAN Server using energy minimized models

#### 3.7.3. Z scores of MUSTER generated models

Ten corresponding template was constructed by MUSTER, but only one template was considered as good (Z score >7.5). The rest had a Z score <7.5, thus was considered as bad (**Table 6**).

**Table 6.** Z scores of different models generated by MUSTER

| Rank | Template | Align length | Coverage | Z-score | Seq_id | Type |
| --- | --- | --- | --- | --- | --- | --- |
| 1 | 1jsuC | 69 | 0.420 | 9.345 | 0.391 | Good |
| 2 | 3jbmA | 153 | 0.932 | 5.296 | 0.124 | Bad |
| 3 | 2mxxA | 155 | 0.945 | 4.759 | 0.116 | Bad |
| 4 | 2muxA | 137 | 0.835 | 4.705 | 0.095 | Bad |
| 5 | 5n5xA2 | 162 | 0.987 | 4.623 | 0.111 | Bad |
| 6 | 4l8nA2 | 160 | 0.975 | 4.564 | 0.106 | Bad |
| 7 | 4fmyA | 162 | 0.987 | 4.426 | 0.086 | Bad |
| 8 | 1su3B3 | 147 | 0.896 | 4.424 | 0.095 | Bad |
| 9 | 4uisA1 | 140 | 0.853 | 4.331 | 0.093 | Bad |
| 10 | 3jb8A | 161 | 0.981 | 4.318 | 0.112 | Bad |

### 3.8. Visualization of selected mutations using Mutation 3D server

Mutation 3D server illustrated selected harmful substitutions (**Figure 7**). Only one domain, CDI (PF02234), of the 164 amino acids in long cyclin dependent kinase inhibitor 1 protein (encoded by CDKN1A gene), was revealed. Among 12 mutations, 3 mutations were uncovered (R19C, Y151C, R156W), the rest 9 mutations (G23D, V25G, V25L, Q29P, F51L, E56K, T57I, G61R, and G61D) were in the domain region, thus considered as higher risk mutation for the cyclin dependent kinase inhibitor 1 protein.

**Figure 7.**
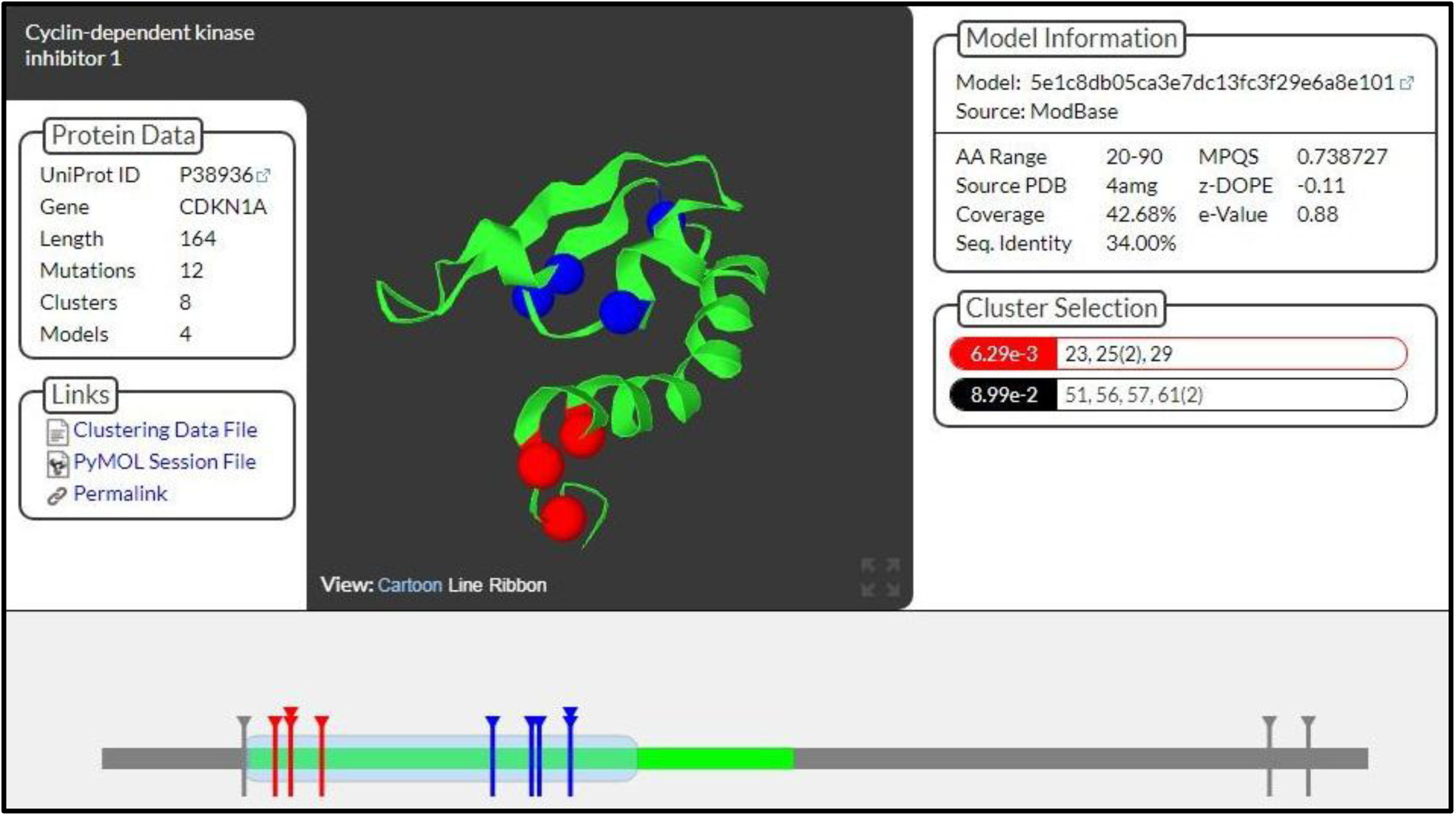
3D structure of cyclin dependent kinase inhibitor 1 protein, generated by mutation 3D server, and representation of selected mutations on protein’s domain

### 3.9. Comparison of total energy and electrostatic constraints between energy minimized native and mutated models

Total energy and electrostatic constraint (expresses as kJ/mol), after energy minimization by GROMOS 96 implementation of Swiss-PdbViewer, of the native (Swiss Model) structure and the mutant modeled structures are shown in **Table 7**. Out of 12 amino acid substitutions, G23D, V25L, F51L, E56K, T57I, G61R, G61D were found to have decrease in both total energy and electrostatic constraint in comparison with the native structure.

**Table 7.**
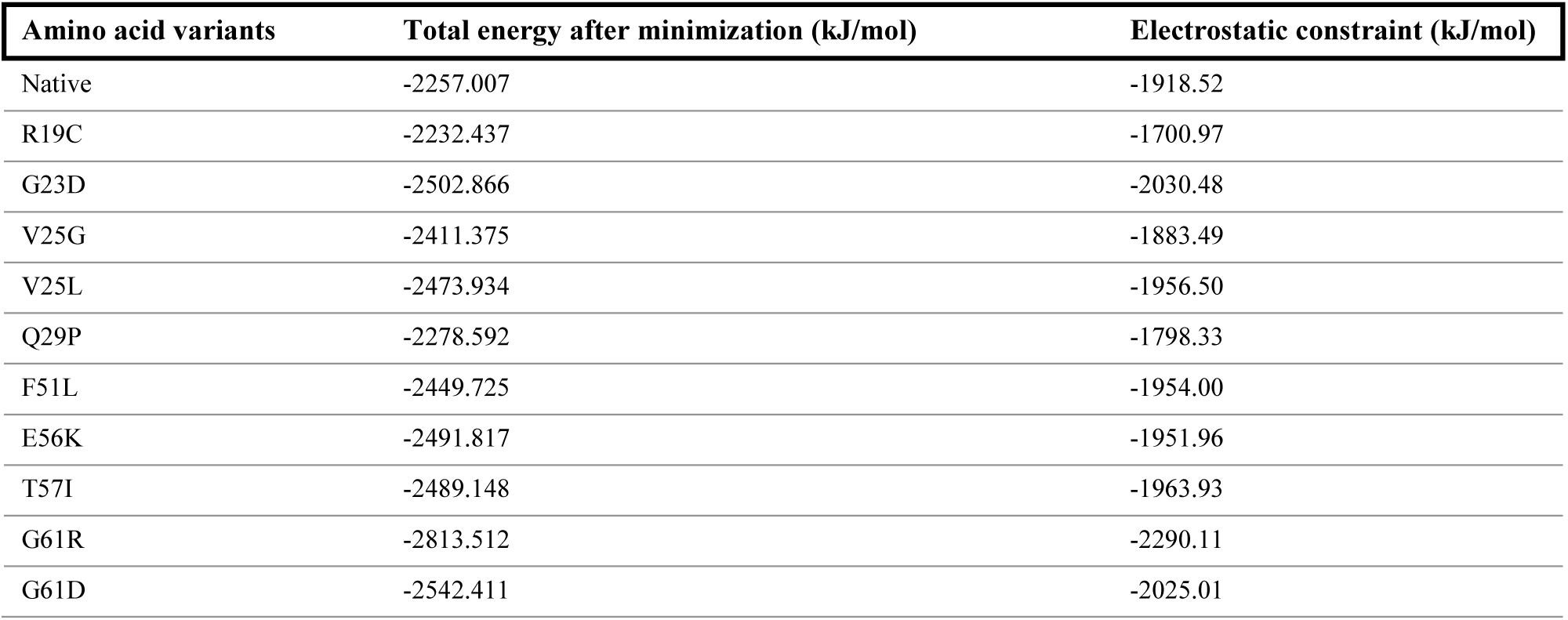
Total energy and electrostatic constraint after energy minimization in native and mutated models

| Amino acid variants | Total energy after minimization (kJ/mol) | Electrostatic constraint (kJ/mol) |
| --- | --- | --- |
| Native | -2257.007 | -1918.52 |
| R19C | -2232.437 | -1700.97 |
| G23D | -2502.866 | -2030.48 |
| V25G | -2411.375 | -1883.49 |
| V25L | -2473.934 | -1956.50 |
| Q29P | -2278.592 | -1798.33 |
| F51L | -2449.725 | -1954.00 |
| E56K | -2491.817 | -1951.96 |
| T57I | -2489.148 | -1963.93 |
| G61R | -2813.512 | -2290.11 |
| G61D | -2542.411 | -2025.01 |

### 3.10. Biophysical analysis of selected amino acid substitutions

Functional effect of missense amino acid substitution, analyzed by Align GVGD, is represented in

**Table 8**. All 12 nsSNPs existed at strongly conserved residues (GV=0). Nine variants, R19C, G23D, V25G, Q29P, T57I, G61R, G61D, Y151C, R156W, were predicted to be in the class C65. Thus, they were inferred as most likely to interfere with the function. One variant, E56K, was predicted to be in class C55. Thus, it was defined as interfering with function. The additional variants, F51L and V25L, were predicted as less likely to interfere with the function.

**Table 8.** Analysis of the functional effect upon nonsynonymous mutation by Align GVGD

| Amino acid variants | A-GVDV |  | Prediction class |
| --- | --- | --- | --- |
|  | GV | GD |  |
| R19C | 0.00 | 179.53 | Class C65 |
| G23D | 0.00 | 93.77 | Class C65 |
| V25G | 0.00 | 108.79 | Class C65 |
| V25L | 0.00 | 31.78 | Class C25 |
| Q29P | 0.00 | 75.14 | Class C65 |
| F51L | 0.00 | 21.82 | Class C15 |
| E56K | 0.00 | 56.87 | Class C55 |
| T57I | 0.00 | 89.28 | Class C65 |
| G61R | 0.00 | 125.13 | Class C65 |
| G61D | 0.00 | 93.77 | Class C65 |
| R156W | 0.00 | 101.29 | Class C65 |
| Y151C | 0.00 | 193.72 | Class C65 |

Classifiers according to A-GVGD, ordered from most likely to interfere with function to least likely:
$GD \geq 65 + \tan(10) \times (GV^{2.5}) \Rightarrow$ Class C65 ⇔ most likely
$GD \geq 55 + \tan(10) \times (GV^{2.0}) \Rightarrow$ Class C55
$GD \geq 45 + \tan(15) \times (GV^{1.7}) \Rightarrow$ Class C45
$GD \geq 35 + \tan(50) \times (GV^{1.1}) \Rightarrow$ Class C35
$GD \geq 25 + \tan(55) \times (GV^{0.95}) \Rightarrow$ Class C25
$GD \geq 15 + \tan(75) \times (GV^{0.6}) \Rightarrow$ Class C15
Else ( $GD < 15 + \tan(75) \times (GV^{0.6})$ ) ⇒ Class C0 ⇔ less likely

### 3.11. Harmful effects of selected variants by MutPred2 server

MutPred2 tool takes the physio-chemical properties in consideration to predict the degree of tolerance for nsSNPs. The summarized results from MutPred server for the selected 12 amino acid substitution is represented in **Table 9**. Gain of intrinsic disorder (P= 0.0065) was predicted at V25G; Amino acid substitution at F51L and E56K was predicted to have a loss of proteolytic cleavage at D52 (P=0.0095); at the substitution G61D, altered metal binding (P=0.0055) was predicted. Amino acid substitution at Y151C was predicted to have loss of phosphorylation at Y151 (P=0.005) along with altered disordered interface (P=0.0019). Loss of acetylation at K161 (P=0.0029) and gain of strand (P=0.0076) were predicted at R156W.

**Table 9.**
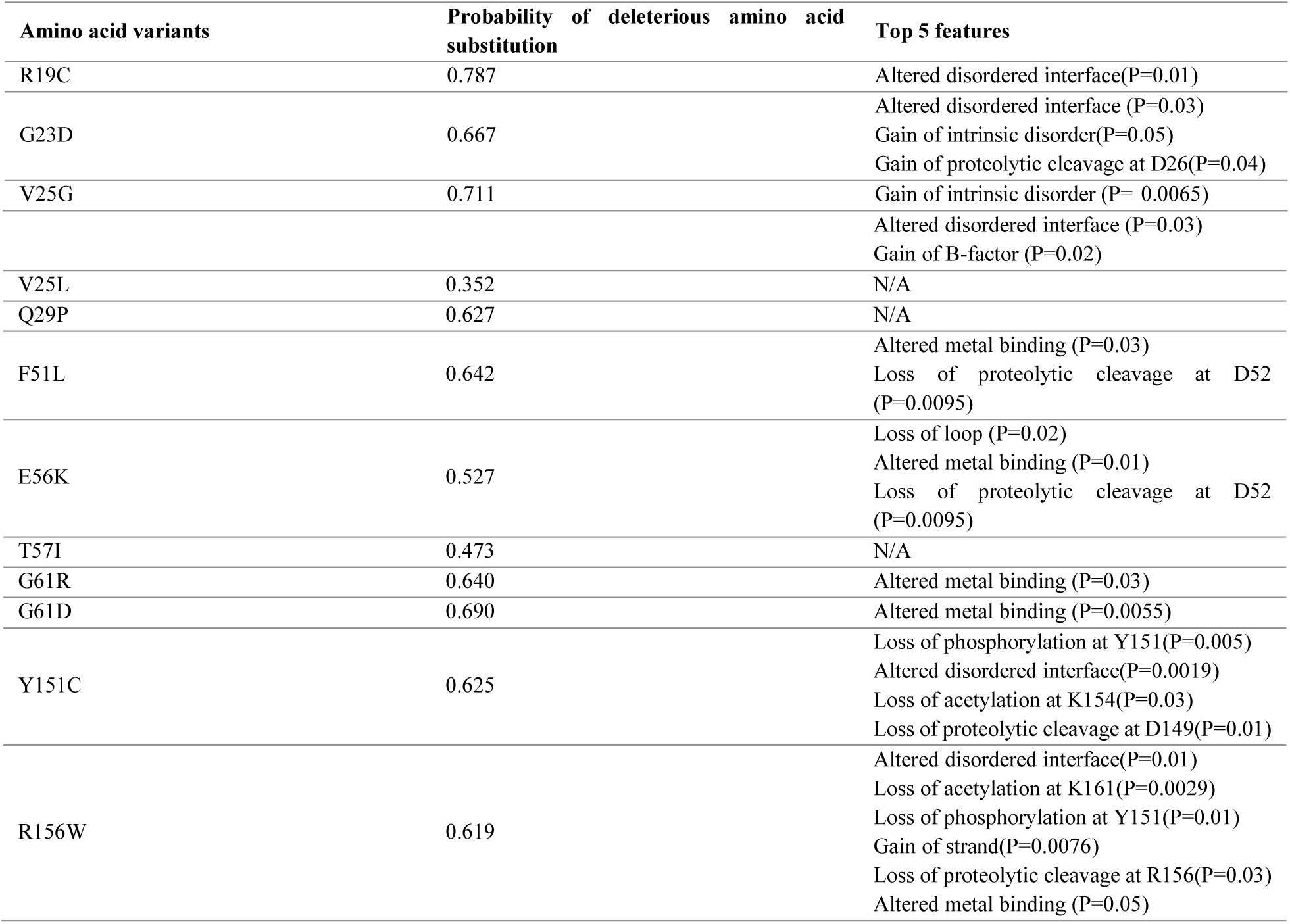
Prediction of the effect of nsSNPs by MutPred server

## 4. Discussion

Non synonymous single nucleotide polymorphism in genes pose a profound impact on the functionality of a protein. The determination of the phenotypic features attributed to nsSNPs by in silico tools may bring the susceptible variants interfering with the natural function. CDKN1A gene, located chromosome 6, codes for the protein p21^Cip1^, also known as cyclin dependent kinase inhibitor 1 protein. This protein inhibits CDK2 and halts cell cycle progression. It acts as a genome guardian by triggering apoptosis upon DNA damage or stress (Georgakilas et al., 2017); this function might become the opposite if mutation in CDKN1A arises (Liu and Kwiatkowski, 2015).

In this in silico study, following the final screening using the SIFT and Polyphen-2 tool, 53 missense substitutions, out of 118 nsSNPs, was predicted to be harmful. In a study to evaluate the effect of missense variants, SIFT, Polyphen-2 and SNAP tool gave better result with higher accuracy than the other to identify deleterious nsSNPs (Thusberg and Vihinen, 2009). Of these 53 nsSNPs of this analytical study, SIFT predicted 43.39% (n=23) substitutions as damaging where Polyphen-2 predicted 64.15% (34) and 52.83% (28) nsSNPs as deleterious according to HumDiv and HumVar scores, respectively. Non synonymous SNPs associated with disease were highest according to MAPP (50), followed by Predict SNP (31), SNAP (29) and PhD SNP (19). Changes in the stability upon mutation was highest according to I-Mutant 3.0 tool (46) and MUpro predicted 41 nsSNPs, out of 53 as responsible to decrease the stability of the protein. I-Mutant 3.0 was found to be the most reliable predictor to identify the changes in stability upon mutation (Khan and Vihinen, 2010). Among these 53 substitutions-R19C (C→T, rs769229553), G23D (A→G, rs368128681), V25G (G→T, rs765248879), V25L (C→G, rs151314631), Q29P (A→C→G, rs756383281), F51L (C→T, rs748842488), E56K (A→G, rs753291170), T57I (C→T, rs772419969), G61R (C→G, rs758877354), G61D (A→G, rs778199042), Y151C (A→G, rs199801759) and R156W (C→G→T, rs374154006) were predicted as both deleterious and responsible for decreasing stability of the protein by all above methods. Thus, further analysis on their effect on the functionality was executed. A similar study based on the dbSNP was conducted to find the deleterious SNPs of BARD1 gene (Alshatwi et al., 2012). Homology model generated by the Swiss-model was taken as native model, considering its overall validation score. Homology modelling has been one of the most promising method to build reliable structures (Bordoli et al., 2008). Following energy minimization, mutant model R19C and V25G presented an increase (less favorable) in the total energy count (kcal/mol) comparing to the native model. Total scores for Y151C and R156W could not be computed due to the limitation in the amino acid length of the model structure (62 amino acid). Only three (R19C, Y151C and R156W) out of these twelve substitutions was uncovered and occurred out of the CDI (PF02234) domain predicted by Mutation 3D algorithm. Domains are the functionally active site of a protein structure and mutations on this site may have a tremendous effect on its activity (Yang et al., 2015). Biophysical analysis by Align-GVGD proposed only two substitutions (V25L and F51L) as less confidently interfering with the function of the protein. More importantly, among these 12 highly damaging mutations-R19C, G23D, F51L, Y151C and R156W occurred at the most conserved site, based on evolutionary history, on the amino acid sequence of CDKN1A, which are predicted to be the positions interfering mostly with the function of the protein. Some molecular features of some selected substitutions revealed by MutPred2 application predicted them as being actionable or confident hypotheses that they might be deleterious or associated with disease. MutPred2 general probability of deleterious mutation score for R19C, G23D, V25G, F51L, E56K, G61R, G61D, Y151C, and R156W was 0.787, 0.667, 0.711, 0.642, 0.527, 0.640, 0.690, 0.625 and 0.619 respectively. The effect of missense mutation at the molecular level was also predicted by in silico analysis in Chinese patients having congenital dyserythropoietic anaemia (CDA) (Wang et al., 2018)

## 5. Conclusion

A simulation based study employed in this study to detect the deleterious nsSNPs using many computational program has revealed, 12 amino acid substitutions (R19C, G23D, V25G, V25L, Q29P, F51L, E56K, T57I, G61R, G61D, Y151C and R156W) of CDKN1A which might be responsible for the functional discrepancies comparing to the novel wild CDKN1A gene having a major role in the regulation of cell cycle progression at G1 phase. Majority of the variants identified in this study are located on domain site and conserved region of cyclin dependent kinase inhibitor 1. In situ evaluation of this gene, and its protein cyclin dependent kinase 1 in this study, which predicted some novel deleterious non-synonymous single nucleotide polymorphisms along with their functional sites and molecular features might come in help for further in vitro study.

